# BABEL enables cross-modality translation between multi-omic profiles at single-cell resolution

**DOI:** 10.1101/2020.11.09.375550

**Authors:** Kevin E. Wu, Kathryn E. Yost, Howard Y. Chang, James Zou

## Abstract

Simultaneous profiling of multi-omic modalities within a single cell is a grand challenge for single-cell biology. While there have been impressive technical innovations demonstrating feasibility – for example generating paired measurements of scRNA-seq and scATAC-seq – wide-spread application of joint profiling is challenging due to the experimental complexity, noise, and cost. Here we introduce BABEL, a deep learning method that translates between the transcriptome and chromatin profiles of a single cell. Leveraging a novel interoperable neural network model, BABEL can generate scRNA-seq directly from a cell’s scATAC-seq, and vice versa. This makes it possible to computationally synthesize paired multi-omic measurements when only one modality is experimentally available. Across several paired scRNA-seq and scATAC-seq datasets in human and mouse, we validate that BABEL accurately translates between these modalities for individual cells. BABEL also generalizes well to new biological contexts not seen during training. For example, starting from scATAC-seq of patient derived basal cell carcinoma (BCC), BABEL generated scRNA-seq that enabled fine-grained classification of complex cell states, despite having never seen BCC data. These predictions are comparable to analyses of the experimental BCC scRNA-seq data. We further show that BABEL can incorporate additional single-cell data modalities, such as CITE-seq, thus enabling translation across chromatin, RNA, and protein. BABEL offers a powerful approach for data exploration and hypothesis generation.

## Introduction

Single-cell technologies have made it possible to precisely characterize cellular state using diverse modalities ranging from gene expression and chromatin accessibility, to proteomics and methylation (Figure 1A)^1^. Such fine-grained understanding provides much more information beyond measuring the bulk average state of a tissue sample, and has enabled novel insights into complex biological systems. However, a notable limitation of standard single-cell technologies is that they only capture one measurement modality (e.g. only RNA-seq or only ATAC-seq) for each cell. This loses the critical information about how the different layers of genomic regulation interact within individual cells.

**Figure 1:**
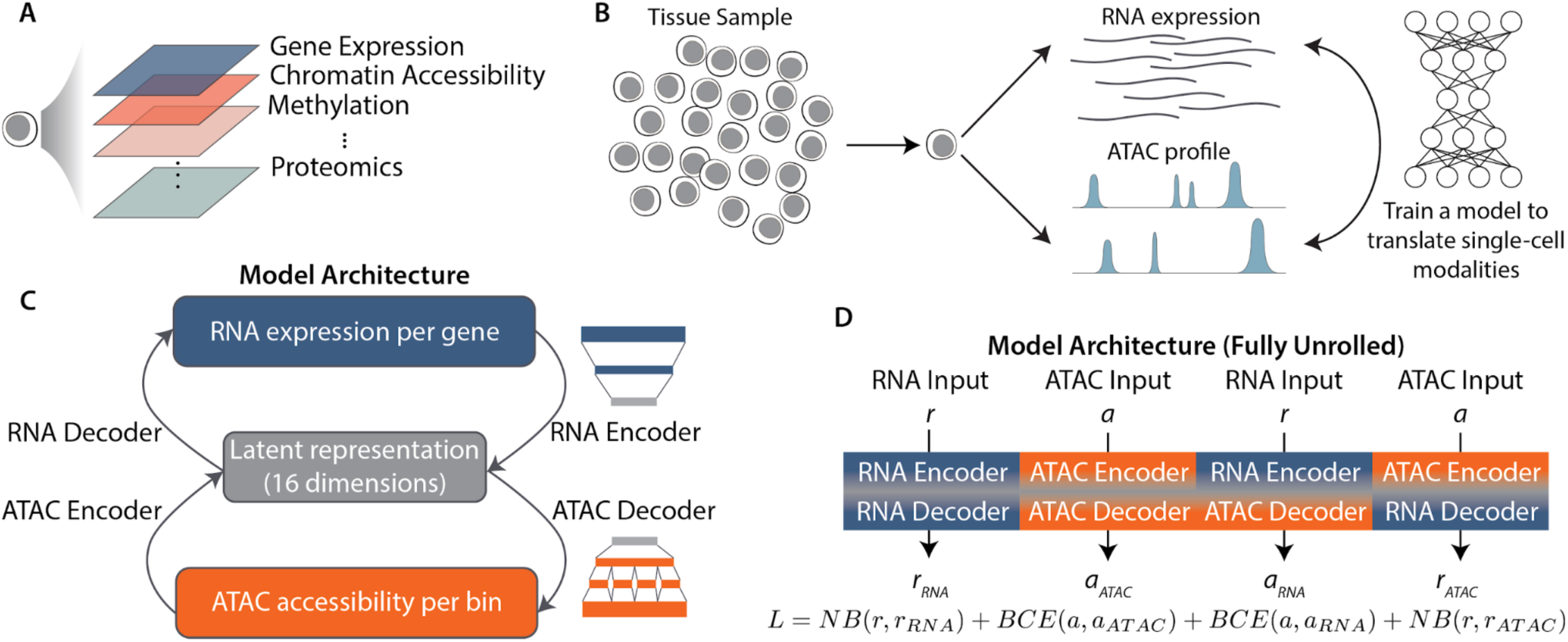
Overview of cross-modality single cell translation with BABEL. (A) Advances in single-cell sequencing technology have enabled a myriad of single-cell modalities, ranging from gene expression to proteomics, as well as technologies jointly measuring combinations of these modalities. However, multi-modal profiling also presents challenges such as increased complexity, noise, and cost. (B) We investigate an alternative approach to single-cell multi-omic profiling by using machine learning to translate between single-cell omics measurements, thus enabling the inference of unmeasured modalities. For our study, we focus on RNA expression and ATAC chromatin accessibility, for which we have the most paired data. (C) Shows BABEL’s modelling strategy; specifically, we have two encoders that project ATAC (orange) and RNA (blue) into a latent space (grey), and two decoders that take points in the latent space and infer the corresponding ATAC or RNA profiles. These encoders and decoders are interoperable by design. The RNA networks use a series of fully connected layers (blue schematic), while the ATAC network breaks into sub-connections that limit the model to learning predominantly intra-chromosomal weights (orange schematic). This greatly reduces model complexity. (D) Summarizes the four possible combinations of the encoders and decoders in our network and shows the joint loss function. We train the model by passing each paired ATAC/RNA observation through every combination of encoders and decoders, and our loss function *L* ensures that all four subnetworks work together to produce accurate translations.

More recently, several multi-omic single-cell methods that jointly profile multiple modalities within the same cell have emerged^1,2^. For example, SNARE-seq and sci-CAR combine chromatin accessibility with RNA gene expression measurements^3,4^, CITE-seq enables joint quantification of RNA expression and protein markers^5^, and Pi-ATAC and ASAP-seq merge epigenomic and protein measurements^6,7^. The paired measurements that these methods generate have helped researchers gain a more comprehensive understanding of how different cellular mechanisms interact. As an example, co-assays of accessibility and expression have been able to specifically identify distal cis-regulatory elements for genes which do not exhibit clear cell type specific promoter accessibility^4^.

However, these joint single cell methods face challenges of their own. Single-cell multi-omics methods often require additional precautions when preserving or isolating cells in order to effectively capture a diverse range of molecules, with RNA often being the most difficult to handle and store^8^. Imperfections in this step can lead to increased noise and drop-out in the resulting data. Seeing as noise and sparsity are already substantial hurdles in analyzing single-cell data^9^, this can make effectively extracting reliable insights from multi-omic single-cell data particularly challenging. Beyond technical feasibility, the increased costs of these multi-omic experiments can also limit the scale at which they can be performed^2^. With these challenges, it may not always be possible or practical to experimentally jointly profile single cells, which motivates the question of how we can extract the most information from samples where only one modality can be captured.

We develop BABEL, a deep learning algorithm that computationally generates, from a single measured modality, other genomic modalities in the same single cell. This enables researchers to perform downstream multi-omic analysis at single cell resolution as if joint profiling data has been collected. This approach is analogous to translating sentences between languages with different grammatical and syntactic structures. BABEL can accurately infer transcriptome-wide single-cell RNA profiles from genome-wide single-cell ATAC profiles, and vice versa (Figure 1B). We focus on RNA and ATAC in this work, because experimental methods for jointly profiling these modalities are more advanced and have the most data available. We expect that BABEL’s approach can be extended to include additional modalities as such data becomes more widely available.

After training BABEL on cells with jointly profiled scATAC-seq and scRNA-seq measurements, we first demonstrate that BABEL performs well on test cell types and tissues that are distinct from the cell population used for training, and where we have paired scRNA-seq and scATAC-seq measurements facilitating comparison. We then show that BABEL can be applied to single-modality single-cell experiments, inferring high-quality cross-domain data that can be analyzed to produce similar conclusions compared to carrying out an entirely separate experiment. We then successfully apply BABEL to analyzing patient basal cell carcinoma (BCC) samples that have been sequenced using single-cell ATAC sequencing (scATAC-seq)^10^. Although this is particularly challenging due to the heterogeneous nature of tumor microenvironments, BABEL’s predictions are concordant with previous findings, and help to uncover additional information compared to previous methods. Throughout our analyses, we find that BABEL’s predictions are consistently driven by individual cell signatures, rather than bulk approximations. Finally, as a proof-of-concept demonstrating BABEL’s versatility, we show that BABEL can predict single cell epitope profiles from scATAC-seq, enabling matched single-cell chromatin, RNA and epitope analysis even though such data is not yet experimentally available.

Several prior works have applied deep learning methods to single-cell data. Many of these focus on developing models to denoise single-cell RNA sequencing (scRNA-seq) data. Examples of these methods include DeepCountAutoencoder, which trains an autoencoder using a (optionally zero-inflated) negative binomial loss to learn a nonlinear function for denoising data^11^, and SAUCIE, which uses a similar autoencoder architecture along with clever regularization techniques to denoise, batch-correct, and cluster scRNA-seq data^12^. Other approaches, such as scVI, apply generative modelling to develop models that can help researchers perform downstream analyses like batch correction and differential gene expression^13^. Single-cell ATAC-seq data has been modeled with machine learning approaches as well, with works like SCALE using autoencoders to learn latent representations conducive to clustering analyses^14^. Deep learning methods for multi-omic data have also been studied, but these prior works did not have access to large-scale paired measurements, which motivated techniques to align latent representations ^15–17^ or constrained these works to bulk measurements^18^. BABEL builds off these prior works while introducing new strategies for more efficient model architectures and latent space learning. To the best of our knowledge, BABEL is the first method to accurately and generalizably translate between gene expression and chromatin accessibility profiles for individual cells.

BABEL addresses a different problem from multi-omic data integration. Prior methods such as iCluster^19^, Seurat^20^, ArchR^21^, MAESTRO^22^, MATCHER^23^, and various matrix factorization approaches^24–26^ excel at data integration, where they take two (typically unpaired) data modalities that have already been measured and compute joint clustering, identify cell-to-cell mappings, and infer cross-domain interactions. BABEL’s goal is to build a generalizable model that takes only one of these modalities and infers the other which can then be used for multi-omic analysis. BABEL provides a powerful tool for hypothesis generation and exploratory analysis, especially in instances where the original sample might no longer be available, cannot be reacquired, or does not contain all necessary molecules, as is often the case for clinical or archival samples^27^.

## Results

### BABEL architecture and design

BABEL consists of four modular neural networks as subcomponents (Figure 1C). Two of these are encoder networks trained to project either RNA or ATAC data into a shared 16-dimensional latent representation. The remaining two neural networks are decoder networks trained to take points in this shared latent representation and infer their corresponding RNA or ATAC profiles. The latent space can be thought of as an abstract representation of cellular state, one that is an intentionally low-dimensional “information bottleneck” to encourage the model to capture major cellular variation, rather than potentially spurious deviations (see Supplemental Note, Supplementary Table 1). Our goal is then to learn encoder models that project single-cell profiles into this cellular state latent space, along with decoder models that can infer observed phenotypes from this latent cellular representation.

While a standard autoencoder maps one data modality onto itself (e.g. scATAC-seq to scATAC-seq), BABEL maps to multiple modalities (e.g. it uses the same scATAC-seq encoder to map to scATAC-seq and to scRNA-seq). This enables the model to be more flexible and efficient. The networks responsible for RNA encoding and decoding consist of fully connected layers that project a continuous, transcriptome-wide expression vector to the latent cellular representation (Figure 1C). The networks responsible for encoding and decoding the binary genome-wide ATAC signal are also composed of fully connected layers, but we leverage the insight that most chromatin accessibility interactions occur at an intra-chromosomal level^28^ to prune the majority of inter-chromosomal connections (Figure 1C). This novel approach substantially reduces the parameter space and helps the model to avoid spurious correlations (see Methods and Supplementary Note for more details).

BABEL is trained using a loss function that requires both encoders to be interoperable with either decoder. This is expressed by enumerating all four possible compositions of our two encoders and decoders and simultaneously training all four of these “paths” through our model to produce correct outputs using four corresponding loss terms (Figure 1D). Under this formulation, the ATAC encoder’s latent output must be consumable by both the ATAC decoder and RNA decoder to produce correct outputs in both output modalities, and the same goes for RNA encoder. This interoperability constraint leverages paired data to alleviate the need for latent space alignment while improving BABEL’s ability to generalize to unseen cell types (see Supplementary Notes, Supplementary Table 2). To evaluate the correctness of inferred RNA values (whether these inferences were generated from an ATAC or RNA input), we use a negative binomial (NB) loss, which has seen success in prior works focused on imputing and denoising single-cell RNA expression^11,13^. To evaluate the correctness of inferred ATAC values, we use a binary cross entropy (BCE) loss – a natural cost function for binary predictions that has been used in prior deep learning models for scATAC-seq data^14^. After training, BABEL can translate between a continuous vector representing an individual cell’s transcriptome-wide RNA gene expression spanning 34,861 genes, and a binary vector of genome-wide ATAC accessibility profiles spanning 223,897 high-resolution peaks with mean and median widths of 796 and 573 base pairs, respectively.

### BABEL performs cross-domain translation with high accuracy

We train BABEL using single-cell multi-omic data jointly profiling ATAC chromatin accessibility and RNA gene expression on the same cells generated on 10x Genomics’ multi-omic platform (see Methods). These data spans cells collected from several human primary cell types and transformed cell lines: peripheral blood mononuclear cells (PBMCs), colon adenocarcinoma COLO-320DM (DM) cells, colorectal adenocarcinoma COLO-320HSR (HSR) cells, and GM12878 cells. We pool and cluster the PBMC, DM, and HSR cells together, reserving one cluster for validation, and one cluster for test, with the remaining cells being used for model training (Figure 2A). Although BABEL itself is agnostic of cluster identity, cluster-based data splits reduce cell similarity between training/validation/test data, challenging the model to generalize to new cell populations. Jointly profiled GM12878 cells are held out from any training purposes, so that they serve as a measure of generalization even more rigorous than the test cluster.

**Figure 2:**
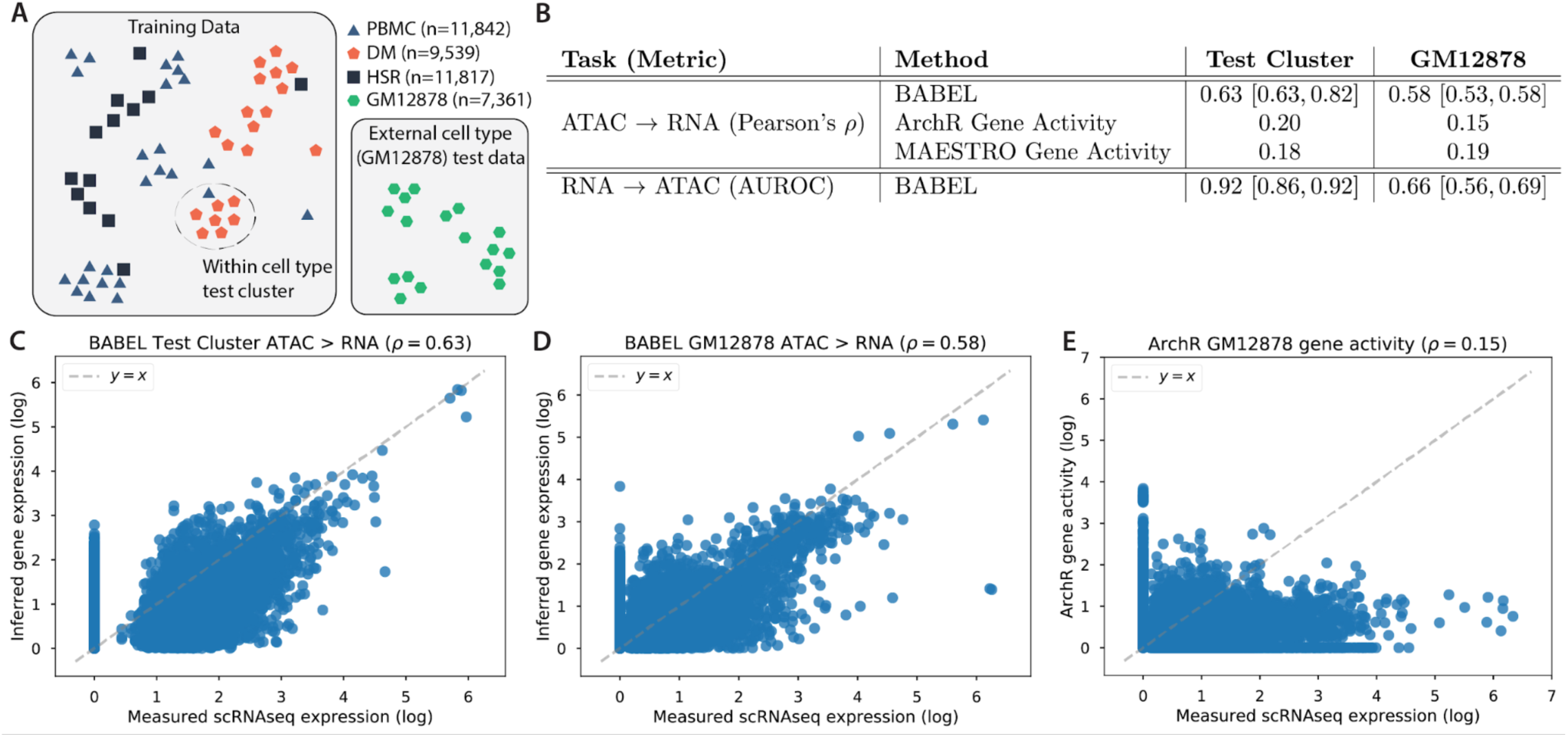
Summary of train and test data, and BABEL’s performance. (A) We train BABEL using a combination of PBMC, DM, and HSR cells with jointly profiled ATAC and RNA measurements. Data from these three are pooled and clustered (“Training Data” box), with one cluster reserved as a test cluster. Splitting by cluster reduces the similarity between train and test data. Moreover, we keep a set of jointly profiled GM12878 cells completely separate from all training purposes, used only for model evaluation (“External cell type” box). (B) BABEL’s performance on the test cluster as well as on the external GM12878 evaluation set, with cross validation performance ranges in brackets. To contextualize BABEL’s ATAC to RNA performance, we include the performance of state-of-the-art gene activity score estimation methods from ArchR and MAESTRO. Plots (C-E) further explore the difference between BABEL and gene activity scores. Each point in (C) represents the expression of one gene in one cell (100,000 such entries randomly chosen for visual clarity) within the held-out test cluster. The x-axis represents the empirically measured expression of that gene in that cell, while the y-axis represents BABEL’s inferred expression. We observe a strong correlation of 0.63 on this test cluster. (D) shows the same plot, but for BABEL’s inferences on the external GM12878 evaluation set instead. (E) shows ArchR gene activity scores generated for the GM12878 cluster, which is less accurate than BABEL.

BABEL achieves strong performance for cross-domain inference on all test data. Inferring RNA expression from ATAC accessibility, it achieves a Pearson’s correlation of 0.63 (Figures 2B, 2C). Inferring ATAC from RNA on this same test set, BABEL achieves an Area Under the Receiver Operating Characteristic (AUROC) of 0.92 (we use different metrics for evaluating ATAC and RNA predictions due to the binary versus continuous nature of these modalities). BABEL’s performance is consistent across cross-validation as well; across five different cluster-based data splits, we observe ATAC to RNA Pearson’s correlations ranging from 0.63 to 0.82 with a median of 0.79, and RNA to ATAC AUROCs ranging from 0.86 to 0.92 with a median of 0.87 (Figure 2B, Supplementary Table 3).

To quantify BABEL’s ability to generalize to a different cell type, we applied BABEL trained on PBMC, DM, and HSR data to paired scATAC-seq and scRNA-seq data from GM12878, without any tuning or modification. This constitutes a challenging test since the lymphoblastoid GM12878 cell line exhibits substantial differences from the three cell types used for training. BABEL generalizes well to the GM12878 external evaluation set as well, with an ATAC to RNA Pearson’s correlation of 0.58 (Figures 2B, 2D), and a RNA to ATAC AUROC of 0.66. BABEL’s performance on the GM12878 evaluation set is also very robust across cross validation folds (where the training and validation sets shift, and we evaluate the five resultant models on the same GM12878 data). The ATAC to RNA Pearson’s correlation ranges from 0.53 to 0.58, whereas the RNA to ATAC AUROC ranges from 0.56 to 0.69 (Supplementary Table 4). Performance is similar for intra-domain translations (i.e. inferring RNA output from RNA input and ATAC from ATAC input), which further validates BABEL’s subcomponent networks (Supplementary Tables 3 and 4).

Since one key objective of BABEL is to infer RNA gene expression from ATAC accessibility, existing tools that infer gene activity scores from ATAC data offer natural benchmarks. Like BABEL, these tools attempt to estimate the gene expression corresponding to an ATAC profile, but take a very different approach – rather than training a model from empirical data, they use hand-crafted formulas built on the intuition that accessibility near a gene should be correlated with the activity or expression of that gene. We specifically evaluated the approaches implemented by ArchR, a state-of-the-art scATAC-seq analysis suite^21^, and MAESTRO, an analysis suite specifically designed for integrative analysis of unpaired scRNA-seq and scATAC-seq data^22^. On our held-out test cluster, the gene activity scores produced by ArchR have a 0.20 Pearson correlation with measured expression, while MAESTRO exhibits a correlation of 0.18 (Figure 2B). These are much lower than the correlation of 0.63 that BABEL achieves. Likewise, evaluating the ArchR and MAESTRO gene activity scores on the GM12878 paired data yields Pearson’s correlations of 0.15 and 0.19, respectively, compared to BABEL’s 0.58 (Figure 2B, 2E).

We additionally investigated how BABEL could perform on non-human data. We trained a separate version of BABEL on paired scRNA-seq/scATAC-seq data from the adult mouse cerebral cortex, generated via the SNARE-seq joint profiling protocol^3^. On the held-out test cluster, BABEL achieves an ATAC to RNA Pearson’s correlation of 0.54, and an RNA to ATAC AUROC of 0.80 (Supplementary Figure 1). Both these values are similar to those we observed on the human dataset, demonstrating that BABEL’s modelling approach is highly generalizable, works across species, and can be successfully trained using data generated by a variety of experimental protocols.

### BABEL’s ATAC to RNA inference captures empirically validated cell-states

It is especially interesting to apply BABEL in settings where we do not have paired measurements, but where BABEL has been trained on reasonably similar cell types. This can enable explorative analyses and hypothesis generation on new data using the computationally imputed paired measurements. As a case study for this application, we reconstruct expression profiles for a set of healthy PBMC cells profiled using scATAC-seq (Supplementary Figure 2A). This data is unpaired (i.e. does not use joint profiling methods) and was generated using a different experimental protocol than was used to generate BABEL’s paired training data, and thus exhibits a different noise pattern that we do not explicitly adjust for. We use the pretrained BABEL to impute transcriptome-wide RNA expression signatures for each cell in the sample. We then apply standard single-cell RNA preprocessing (i.e. size normalization followed by log-transformation) to the inferred expression, and visualize the cells using the Uniform Manifold Approximation and Projection (UMAP) algorithm^29,30^ applied to their imputed RNA expression signatures. We color each cell by its original ATAC-based cell type (Supplementary Figure 2A) to visualize how well these are retained in BABEL’s imputed RNA profiles (Figure 3A).

**Figure 3:**
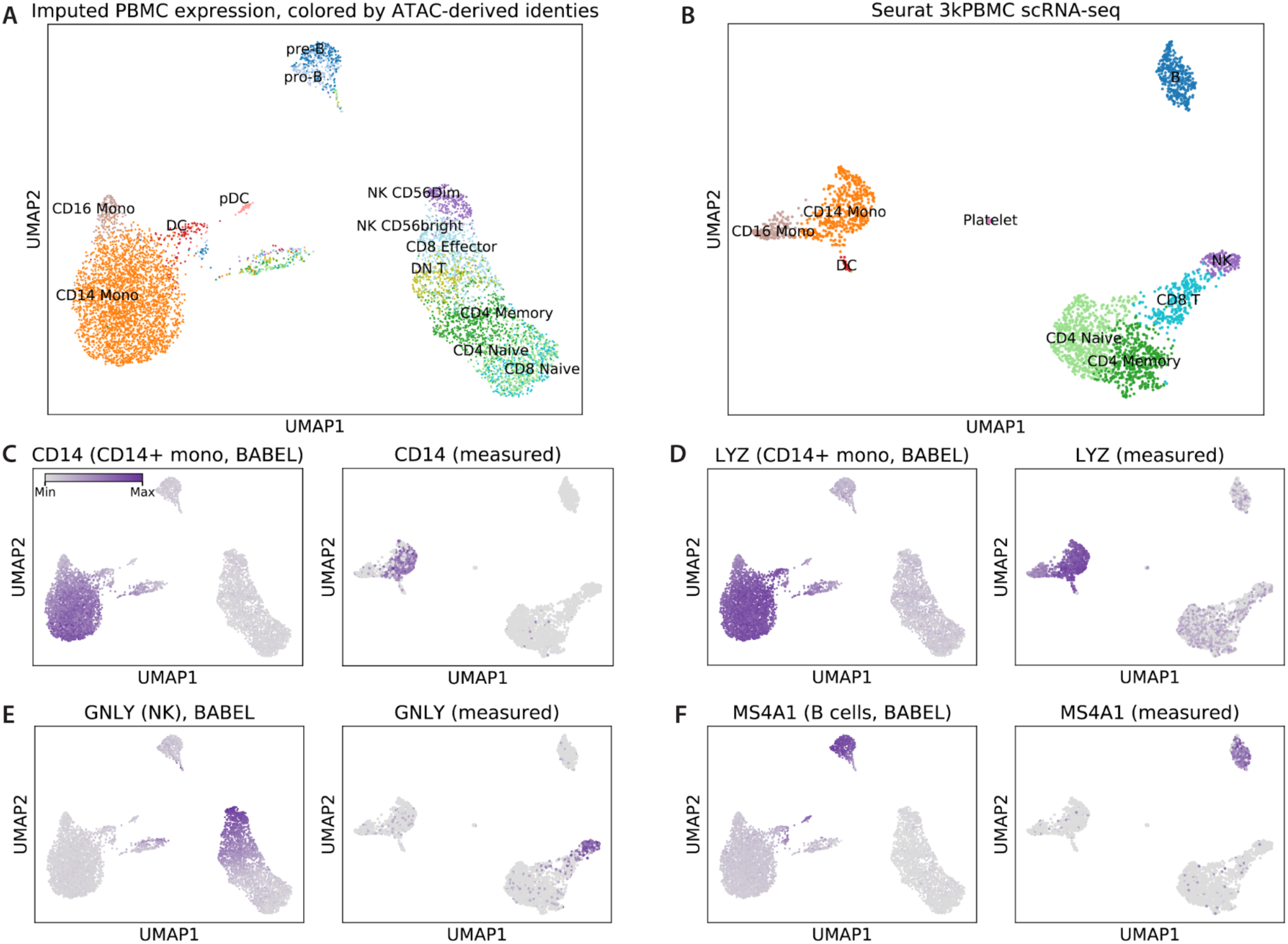
BABEL’s ATAC to RNA translation closely matches empirical results on unpaired PBMCs. (A) UMAP visualization of single-cell expression profiles imputed by BABEL from scATAC-seq, colored by ATAC-derived cell type identities. (B) UMAP visualization and cell types in empirically measured PBMC scATAC-seq for comparison. These two plots exhibit highly concordant global structure (i.e. both show three main cell groups), as well as very similar inter-cell type relationships. (C) Highlights the expression of *CD14* (a marker for CD14+ monocytes) within the inferred gene expression data (left) and the empirically measured data (right). In both cases, *CD14* expression is highly correlated with CD14+ monocytes, as expected. (D) shows the same plots, but highlighting *LYZ*, which is also a marker for CD14+ monocytes. (E) and (F) highlight expressions of *GNLY* and *HS4A1*, which are markers for natural killer and B cells, respectively.

For comparison, we include a parallel tissue-matched analysis of single-cell RNA expression in healthy PBMCs, where we use Seurat to cluster and visualize these cells (Figure 3B). We find many similarities between our imputed UMAP visualization compared to this empirical visualization. There are three major clusters in both the imputed and empirical data. One major cluster corresponds to B-cells (Figure 3A, 3B, top region), one primarily consists of CD14+/CD16+ monocytes and dendritic cells (left region), and one primarily contains CD4, CD8, and natural killer (NK) cells (bottom right region). These results suggest that the imputed single-cell gene expression retains much of the global gene expression patterns and relationships that are empirically observed. Further analysis also reveals that this overall concordance is consistent across different versions of BABEL trained on different cross-validation folds (Supplementary Table 5) and is also recapitulated when we apply UMAP visualization to BABEL’s shared latent representation (Supplementary Figure 2B). This consistency suggests that BABEL can recognize complex relationships between cells in its input and leverages these biological relationships when generating latent representations and when predicting transcriptomic profiles. We believe this property helps BABEL generalize to more contexts (see Supplementary Note) and suggests that BABEL’s latent representation could be a potentially interesting basis for downstream analyses like clustering or lineage tracing.

In addition to examining cell cluster concordance, we further evaluate how well BABEL imputes expression of certain well-known marker genes correlated with specific cell types. *CD14* is a canonical marker for CD14+ monocytes. By coloring each cell in the imputed gene expression UMAP visualization by its imputed *CD14* expression, we find that *CD14* expression coincides very well with CD14 monocyte cells (as identified via standard scATAC analysis methods) (Figure 3C, left panel). As expected, the experimentally measured *CD14* expression overlaps nearly perfectly with CD14+ cells as well (Figure 3C, right panel for UMAP of experimental scRNA-seq). Performing a similar comparison for *LYZ*, another well-known marker for CD14+ cells, we see that BABEL likewise reproduces experimentally validated expression distributions (Figure 3D). Such concordance extends to other cell types as well. Examining the expression of *GNLY*, a marker for natural killer cells (Figure 3E), and *MS4A1*, which corresponds to B cells (Figure 3F), we consistently see that the imputed expression of marker genes matches the cell-types they correspond to, just as the empirically measured data does. This shows that BABEL is not just predicting an average expression of every gene across all cells regardless of the ATAC profile, and that it is capable of highly specific expression inference for individual cells. These results suggest that BABEL can facilitate down-stream cell type analysis by computationally generating missing data modalities. Although BABEL’s strong performance here benefits from having seen similar PBMC cells in its training set (see Supplementary Note, Supplementary Figure 3), generalizing to this external PBMC dataset is still challenging as it was generated by different experimental protocols than those used to generate BABEL’s training data.

### BABEL can provide new insights for patient samples

We next apply BABEL to data acquired from basal cell carcinoma (BCC) tumors to investigate its application on challenging patient samples without any additional fine tuning or training. These samples, acquired from seven BCC patients, represent malignant, stromal, and immune cells present within the tumor microenvironment (TME) both before and after anti-programmed cell death protein 1 (PD-1) immunotherapy (PD-1 blockade), and were profiled using scATAC-seq to generate chromatin accessibility profiles for 37,818 cells^10^. These cells were originally analyzed using standard scATAC-seq methods by calculating gene activity scores using Cicero^31^, and using these scores to label cell types and infer lineages.

A superset of these BCC samples have also been studied in a separate, unpaired experiment using scRNA-seq profiling^32^, which offers an opportunity to evaluate aggregate concordance metrics. Specifically, we leverage the intuition that by averaging across all cells in a single cell experiment, the resulting pseudo-bulk expression values should be comparable across related experiments even in the absence of paired measurements. We generated pseudo-bulk RNA expression profiles from the single-cell RNA expression BABEL inferred from scATAC-seq, compared these against the empirical pseudo-bulk generated from the corresponding patients’ tissue-matched scRNA-seq experiment, and found good agreement (Pearson’s ρ = 0.70, Figure 4A). This concordance also holds if we examine specific cell types, instead of the global population (Supplementary Figure 4). Furthermore, the mismatch between BABEL prediction and empirical scRNA-seq tended to concentrate among genes with low observed expression which could have been under-reported experimentally due to drop outs. For context, we also calculated the pseudo-bulk correlation of Cicero’s gene activity scores against the empirical pseudo-bulk, and observed a weaker correlation of 0.27 (Figure 4B). Overall, BABEL’s predicted RNA expression profiles based on BCC scATAC-seq are robust, though somewhat less so than on PBMCs (see Supplementary Notes). Therefore, rather than applying UMAP to visualize these cells based on their predicted RNA signatures as we did for the PBMCs, we simply visualize the predicted expression with respect to the ATAC-based UMAP projection for these cells (Figure 4C). ρ.

**Figure 4:**
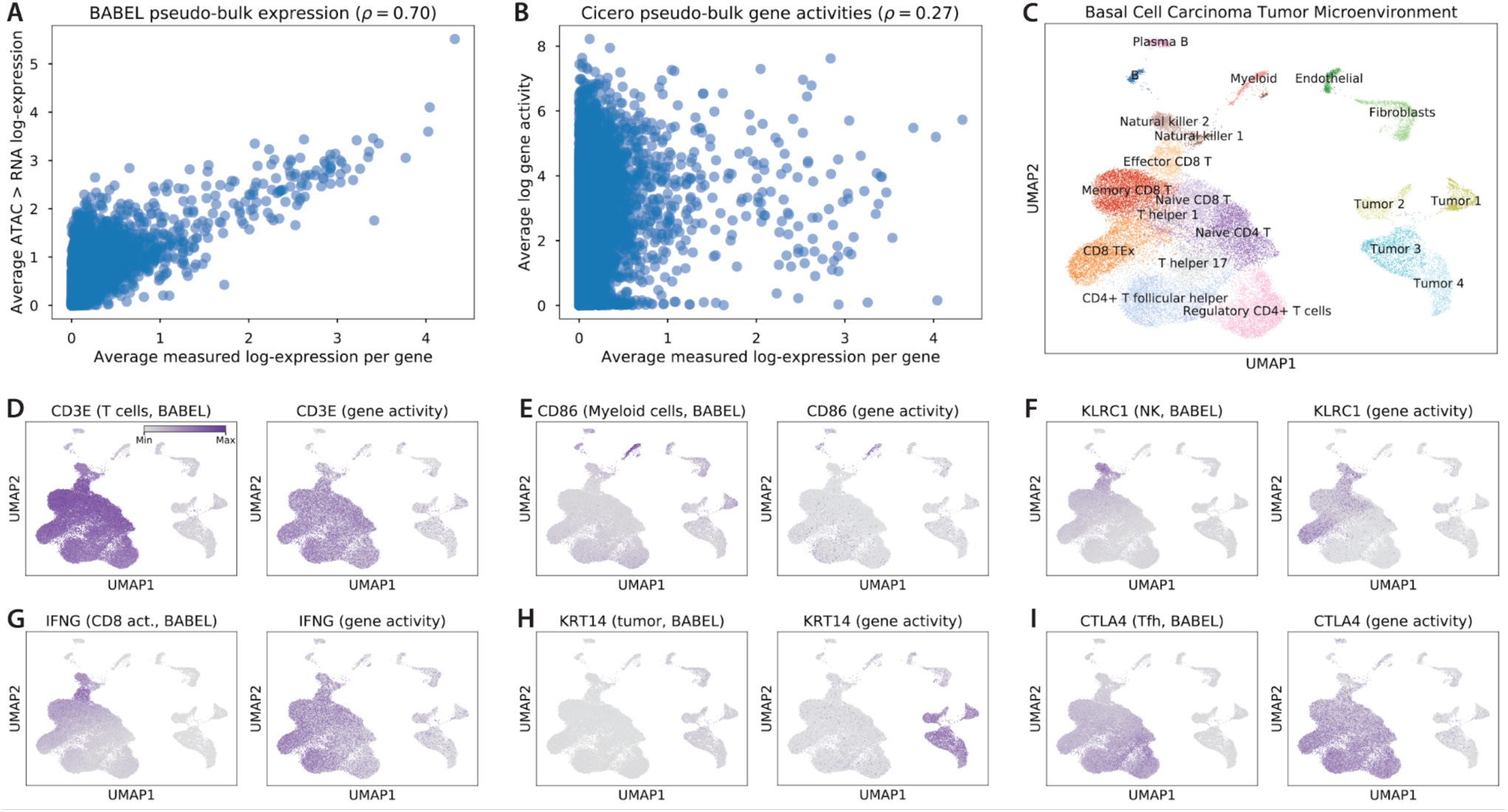
BABEL makes accurate predictions on clinical basal cell carcinoma (BCC) scATAC-seq samples and generates new interpretations. (A) Correlation of transcriptome-wide pseudo-bulk expression: x-axis represents expression within a tissue-matched scRNA-seq study averaged across cells, y-axis represents similarly averaged BABEL predicted single-cell expression from scATAC-seq data. The strong correlation, especially compared to a similar plot for gene activity scores (B), suggests that BABEL generalizes well to patient cancer samples. (C) ATAC-based UMAP visualization of BCC TME cells (reproduced with permission)^10^. We use this projection as a scaffold to visualize imputed single-cell RNA expression. (D) Highlights the imputed expression of *CD3E* (left), a T cell marker, compared to its corresponding inferred gene activity score (right). BABEL recapitulates expected expression of *CD3E* here, as well as for the myeloid marker *CD86* (E). For natural killer marker *KLRC1* (F) and CD8 activated T cell marker *IFNG* (G), BABEL predicts more localized, specific expression than gene activity scores. However, for genes that have little to no presence in the training data such as *KRT14*, BABEL shows weaker performance (H). BABEL also lets us expand on prior conclusions, predicting more distinct overexpression of immunosuppressive genes like *CTLA4* (I) in the Tfh cluster, which strengthens these cells’ reported similarity to exhausted CD8 cells.

The original scATAC-seq BCC report identified several populations of CD4^+^ and CD8^+^ T cells, along with B cells, natural killer (NK) cells, malignant tumor cells, among others (Figure 4C). We next explored whether BABEL could accurately predict cell type specific expression for marker genes corresponding to each cell type. *CD3E* is a marker gene known to be expressed in T cells, and by overlaying predicted expression values upon the scATAC-seq UMAP projection, we see that BABEL recovers this relationship (Figure 4D, left panel), much as gene activity scores do (Figure 4D, right panel). *CD86*, a marker for myeloid cells, exhibits high specificity and concordance with gene activity scores as well, with predicted expression highly specific to labelled myeloid cells (Figure 4E). Predicted single-cell expression of *KLRC1* (Killer Cell Lectin Like Receptor C1), a marker for NK cells, is also highly concordant with both annotated cell types and gene activity scores (Figure 4F). In fact, the distribution of BABEL’s imputed *KLRC1* expression appears to be even more specific to the NK cell clusters than the gene activity scores – Cicero gene scores also predict *KLRC1* activity in epigenetically similar cytotoxic CD8^+^ T cell clusters^33,34^. We observe similar specificity improvements for *IFNG*, which encodes the interferon gamma protein and is specifically expressed in CD8^+^ activated T cells in scRNA-seq experiments measuring these BCC samples^32^. In this context, BABEL’s predicted *IFNG* expression is much more localized than the corresponding *IFNG* gene activity scores (Figure 4G), and is thus much closer to empirical scRNA-seq for these cells. These two examples highlight concrete cases where BABEL offers clear improvements over traditional gene activity scores. However, BABEL is less accurate if the cell type corresponding to a marker gene is not present in the training data. For example, *KRT14* is a prominent marker for malignant BCC cells, but since there are no basal epithelial cells in our training data, BABEL predicts a relatively weak signal for *KRT14* (Figure 4H).

BABEL can also help develop more nuanced understandings for certain cell types in this study. The original scATAC-seq BCC study identified a set of T follicular helper (Tfh) cells that exhibited striking epigenomic similarity to exhausted CD8 T cells (TEx), suggesting that Tfh and TEx differentiation may be driven by a shared regulatory program^10^. Using BABEL’s predictions, we find that this Tfh cell cluster may also overexpress several common immunosuppressive genes, such as *CTLA4* (Figure 4I), *PAK2*, and *FAS*^35–37^. While *CTLA4* is also an overexpressed marker for these cells based on gene activity scores, *PAK2* and *FAS* gene activity scores exhibit weak to no overexpression, despite all three of these genes being overexpressed in Tfh cells in the tissue-matched scRNA-seq study^32^ (Supplementary Figure 5). Here, BABEL’s ATAC-based inferences help recover an additional parallel between TEx and Tfh cells that is confirmed by scRNA-seq, whereby these cell types not only share epigenetic similarity, but also are also similar in their overexpression of several immunosuppressive factors.

### BABEL can be extended with additional data modalities

We finally demonstrate that BABEL can be easily extended to predict additional data modalities, such as protein epitope profiles. We trained an auxiliary protein epitope decoder network using 33,455 jointly-profiled RNA and protein epitope measurements of human bone marrow cells, measured using CITE-seq^20^. This protein decoder network leverages BABEL’s pre-trained latent representation as a starting point for inferring protein epitope profiles (see Methods for more details). The resulting protein decoder network combines with BABEL’s RNA encoder to accurately impute epitope profiles from scRNA-seq on test bone marrow cells (Supplementary Figure 6A) and on new PBMC cells measured by CITE-seq (Supplementary Figure 6B). As a proof-of-concept, we show that BABEL’s interoperable structure also enables us to generate epitope profiles of a cell from its scATAC-seq data (Supplementary Figures 6C-E). The fact that BABEL was able to make these predictions without training on the ATAC-protein modality pair further illustrates the potential power and versatility of its computational cross-modality translation approach. As more single-cell modalities become available, BABEL can flexibly incorporate these data and link increasingly diverse layers of molecular information, even across modalities that have yet to be jointly profiled.

## Discussion

BABEL learns a set of neural networks that project single-cell multi-omic modalities into a shared latent representation capturing cellular state, and subsequently uses that latent representation to infer observable genome-wide phenotypes. To achieve our specific task of translating between RNA gene expression and ATAC chromatin accessibility profiles, we leverage the best practices for modelling scRNA-seq described in previous works, while introducing two novel techniques. First, BABEL’s encoder and decoder networks for ATAC data are designed to focus on more biologically relevant intra-chromosomal patterns. Second, BABEL’s interoperable encoder/decoder modules effectively leverage paired measurements to learn a meaningful shared latent representation without the use of additional manifold alignment methods. We demonstrate that our resultant BABEL model performs well across a variety of contexts, including held-out test clusters, data generated from different experimental protocols, and even data generated from aberrant patient carcinoma samples involving different tissues from those used to train BABEL. While our evaluation focuses on predicting gene expression from accessibility due to the relative interpretability of gene expression, we expect that the opposite expression-to-accessibility translation could be useful in other contexts. By providing paired RNA gene expression and ATAC chromatin accessibility single cell measurements without costly experiments, BABEL can be a valuable tool for hypothesis generation and exploration.

We also carefully investigated potential limitations of BABEL. Across our experiments with unpaired PBMC cells and BCC patient samples, BABEL performs best when it is asked to make predictions on cells for which it has seen similar training examples. This limitation is shared by most machine learning approaches; samples that deviate too far from the training set often exhibit poor predictive performance^38^. As a practical metric, we suggest that aggregate pseudo-bulk correlation can be a good measure of the quality of BABEL’s imputation on new data. Researchers using BABEL could easily compute such pseudo-bulk correlations against a growing library of publicly available (bulk or single cell) reference datasets to evaluate whether BABEL produces trustworthy inferences for their specific experiments.

Beyond translating between RNA and ATAC, we also demonstrated that BABEL provides a computational framework that can be extended to translate between other single-cell modalities by adding additional encoder and decoder networks. Generalizing BABEL to additional modalities or to new tissues would require additional joint single-cell profiling data. Given these measurements, BABEL can act as a pretrained network for transfer learning, especially since its components are interoperable. This can reduce the amount of new data required. New modalities could learn to project into or predict from the same pre-defined latent space, which simplifies the learning problem and enables translation between unmeasured domain pairs, as shown with BABEL’s ATAC to protein predictions.

BABEL can be combined with other analysis workflows to improve multi-omic integration. For example, single-cell gene integration algorithms commonly rely on gene activity scores to bridge the gap between ATAC and RNA modalities, thus allowing cluster-to-cluster or even cell-to-cell mapping/integration between separate ATAC and RNA experiments; substituting gene activities for BABEL’s more accurate RNA expression inferences could greatly improve these integration techniques. Another way that BABEL could be extended is by integrating its multi-omic approach with existing unimodal models that use variational inference techniques to perform clustering, batch correction, or other similar tasks, thus enabling a new class of multimodal algorithms. BABEL’s ability to impute unseen modalities can also be combined with multi-view clustering to more robustly identify groups of related cells.

More broadly, we envision BABEL as a powerful tool for single-cell regulome analysis in an increasingly multi-omic world. As methods for measuring different modalities of information within a cell become more available, we will increasingly face tradeoffs between the number of modalities, depth of analysis, sample number, and cost. Approaches like BABEL can greatly improve data efficiency beyond the Pareto frontier defined by technological limitations of co-measurement. Once a class of samples has been jointly profiled by single-cell multi-omic approaches, scientists can study future instances of such samples (in detailed time courses, perturbation, etc.) with the most economical or technically feasible modality, and infer the remaining information using BABEL. These considerations may be particularly valuable for human clinical samples, which are limited in quantity and perhaps stored in archival formats that do not permit all modalities to be measured^27^. By analogy, in 1804, Lewis and Clark took two years to explore, map, and journey from St. Louis to the Pacific coast, but travelers today can easily navigate this route in a matter of days by leveraging existing information rather than painstakingly remapping the terrain each time. We hope that BABEL, along with other multi-omic data inference tools, may provide similarly rapid and cost-effective data navigation and insights going forward.

## Methods

### Data preprocessing

We consider scRNA-seq data as continuous values. To preprocess the scRNA-seq data, we start with a matrix of unnormalized counts per gene per cell generated using Hg38 (or mm10 for SNAREseq mouse data). These can be produced using tools such as CellRanger. We first perform basic filtering, removing genes that are encoded on sex chromosomes and removing cells that express fewer than 200 genes, or more than 7000 genes (2500 for SNAREseq mouse data). We then size-normalize the data, such that each cell’s total counts sums to the median number of counts per cell. We then log-transform the size-normalized counts and standardize these to have zero mean and unit variance. We also clip values within the top and bottom 0.5% of the overall distribution. When preprocessing external scRNA-seq data for model evaluation (i.e. not training), we perform this same series of preprocessing steps.

We regard scATAC-seq data as binary values, as we found that regarding scATAC-seq data as continuous made the prediction problem significantly more difficult, without adding to the usefulness of those inferences (i.e. not providing a meaningful measure of increased accessibility). To preprocess scATAC-seq data, we start with a cells by peaks matrix generated using Hg38 (or mm10 for SNAREseq mouse data). Such matrices can be produced by tools like ArchR or Signac. From there, we remove peaks derived from sex chromosomes, merge overlapping peaks, and binarize the data by replacing all non-zero values with a value of 1. We then perform a few basic filtering steps, removing peaks that occur in fewer than 5 cells or more than 10% of cells. Removing overly rare peaks helps prevent the model from overfitting on just a handful of examples, while removing overly common peaks helps the model focus on learning important variation between cells. Overall, the ATAC input that our model sees is a filtered, binarized view of the original peaks.

When assessing the model’s generalizability on ATAC inputs that may not exactly match the specific peaks that BABEL is trained on, we re-pool the input ATAC peaks to match BABEL’s peaks. This re-pooling process involves first using a liftOver tool (available from https://genome.ucsc.edu/index.html) to convert Hg19 coordinates to Hg38 coordinates, if appropriate. We then take each source/input peak, determine which BABEL peak(s) it overlaps, and transfer the source peaks’ values to the overlapped peaks(s) using an “or” operator to combine/reduce multiple values. In cases where a source peak has no such overlap, its value is dropped. As a concrete example, if one of the model’s learned ATAC peaks was chr1:1000-2000, and it was asked to evaluate an input with peaks chr1:950-1150 and chr1:1190-2090, the input to the model’s chr:1000-2000 peak would be the result of taking a “or” operator of the values at the two input peaks. This process can reduce the resolution of input datasets, but we find that BABEL is able to make robust predictions regardless.

Many of the described preprocessing steps are done via the Python software packages Scanpy version 1.4.3 and AnnData version 0.6.22^39^.

### Data splits

Training, validation, and test splits were defined by clustering the scRNA-seq data. Specifically, we apply the Leiden clustering algorithm^40^ with a resolution of 1.5 on size-normalized, log-normalized counts (see Data preprocessing section). Based on the clustering results, we assign the two largest clusters to be the validation and test set cell clusters. The cells in the remaining clusters make up the training set. By creating training/validation/test splits by clusters rather than randomly, we reduce the model’s propensity to perform well on validation or test sets simply by “memorizing” a similar cell that it has seen before in training. Similarly, performance on the held-out test cluster is thus a stronger indicator for how well the model will generalize.

For cross-validation, we evaluate five different combinations of validation and test clusters; no cluster is used more than once as a validation set, or more than once as a test set. These splits are defined by rotating through the five largest clusters. The first cross validation split corresponds to using the aforementioned two largest clusters as validation and test, and is the primary model we use to report non-cross-validation results in our manuscript. As is customary, we use the validation set for hyperparameter and architecture tuning, as well as for early stopping.

### BABEL architecture

BABEL was implemented using a combination of the PyTorch (version 1.2.0) and Skorch (version 0.7.0) libraries. BABEL consists of four primary components: two encoder networks, and two decoder networks. Each encoder is responsible for projecting either RNA or ATAC input into a shared latent space, and each of our two decoders is likewise responsible for projecting from the shared latent space to inferred RNA or ATAC outputs. These networks are designed to be interoperable – the RNA encoder is compatible with both the RNA decoder and the ATAC decoder, and similarly for the ATAC encoder. Each decoder is also interoperable with both encoders. Note that BABEL does not leverage cluster information in its modelling approach; by not conditioning predictions on cluster identities, which may shift between datasets, BABEL can more easily generalize to novel datasets.

The RNA decoder outputs two parameters, mean and dispersion. The mean represents an estimate of the “denoised” expression, and the mean and dispersion jointly describe the likelihood of the observed expression values under a negative binomial distribution. Existing works that train models on scRNA-seq data like DeepCountAutoencoder^11^ and scVI^13^ use a similar parameterization for their RNA outputs. Coupled with a negative binomial loss, this helps models learn to infer the true, “de-noised” RNA expression values rather than the noisy observed values. The ATAC output, on the other hand, consists of a single output bounded between [0, 1]. These ATAC outputs can then be binarized to obtain a more conventional ATAC profile using an approach developed for SCALE^14^, where each entry in the predicted cell by peak matrix is set to “1” if its value is greater than both the corresponding column and row means and “0” otherwise.

For the RNA encoder, we project the input vector of genome-wide expression (n=34,861 for the human model, n=22,541 for the mouse model) to 64 dimensions, followed by a Parametric ReLU (PReLU) nonlinear activation. We then project to a 16 dimensional shared latent space, again with a PReLU nonlinearity. The RNA decoder can be thought of as the “inverted” version of this network architecture. We start by applying a fully connected layer and a PReLU activation to project the 16 dimensional shared latent representation to a 64 dimensional hidden layer. We then take this project this 64 dimensional layer into two different outputs of size N. The first output is passed through an exponential activation, and the second is passed through a softplus activation; these two outputs thus estimate the mean and dispersion parameters describing a per-gene negative binomial distribution, respectively.

For the ATAC encoder and decoder, we leverage the insight that most DNA accessibility interactions occur on an intrachromosomal level, rather than an interchromosomal level^28^. When designing the ATAC encoder, instead of using a fully connected network that connects the entire set of input ATAC peaks spanning the entire genome to a hidden layer, we instead use a series of much smaller fully connected layers, each connecting a single chromosome’s ATAC peaks to a subset of the next hidden layer. We apply this twice, such that each chromosome’s ATAC peaks are transformed into a 32 dimensional representation (with PReLU activation), followed by a 16 dimensional representation (with PReLU activation), which is then concatenated across all chromosomes. For the human genome with *C*=22 autosomes, this concatenated layer would have 16 x 22 = 352 dimensions. We then finally project this concatenated layer to our 16 dimensional shared latent representation, again using the PReLU activation. As this final layer is not split by chromosomes, it allows the model to learn some inter-chromosomal interactions. The ATAC decoder can be thought of as the “reversed” version of this ATAC encoder network as well. We start by projecting the 16 dimensional latent representation to a *C* x 16 dimensional hidden layer (with PReLU activation), which is then split into *C* blocks of 16 dimensions each. Each of these *C* blocks then passes through its own set of fully connected layers and PReLU activations to map into a hidden layer of 32 dimensions, and then to the outputted, per-chromosome ATAC peak predictions with a final sigmoid activation function that bounds the output values between 0 and 1. Each chromosome need not have the same number of output/input peaks. For comparison, a naive approach where all ATAC connections are genome-wide would necessitate nearly *C* times as many parameters (see Supplementary Note).

### BABEL training

As previously mentioned, the RNA decoder outputs a mean *ŷ* and dispersion *θ* vector for each gene. Together, these two vectors parameterize the likelihood of observing the measured scRNA-seq expression *y* on a per-gene basis under a negative binomial (NB) distribution, as shown below (Г denotes the gamma function).

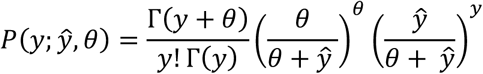

Intuitively, we want to find the mean and dispersion parameters *ŷ, θ* that maximize the likelihood of the observed data. This is equivalent to minimizing the negative log likelihood, which we use as our loss as follows (*ε* is a small constant we introduce for numerical stability).

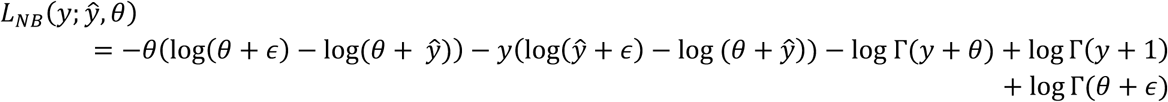

Since the ATAC decoder is tasked with a binary prediction task for each peak across the genome, we use a binary cross-entropy (BCE) loss where *x* represents the measured binary ATAC signal at that bin, and 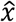 represents our model’s corresponding prediction:

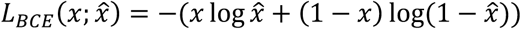

Given these building blocks, we can then build the overall loss function for our model. Recall that we want each encoder to be composable with either decoder, such all our networks are interoperable. We express this in our overall loss function *L* by including a term for each of our four encoder/decoder combinations. Here, *r* and *a* denote the measured, “true” RNA and ATAC signals, respectively, and subscripts denote the source modality that was used to infer either RNA or ATAC (e.g. *r*_*ATAC*_ represents the inferred RNA values from an ATAC input).

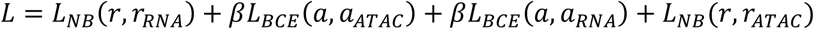

The first two terms encapsulate how well our model can “reconstruct” the RNA and ATAC inputs by performing basic intra-domain inference. The latter two terms encapsulate how well our model does cross-domain prediction (i.e. inferring ATAC profiles from RNA expression and vice versa). The constant term *β* ensures that the binary cross-entropy and negative binomial losses are numerically within the same order of magnitude. Based on manually examining the magnitude of NB and BCE losses over the first few training epochs, we set *β =* 1.33 for all results described in this paper.

We train our model using the Adam optimizer^41^ with a batch size of 512 and a learning rate of 0.01. We reduce the learning rate when validation set loss plateaus, along with early stopping based on validation set loss. We also use batch normalization and gradient clipping to aid in training stability.

While other works that learn a shared latent space between multiple data modalities tend to do so by using an adversarial network in the latent space to distinguish between latent points that do or do not match the latent space distribution^17^, we find that this is unnecessary (see Supplementary Notes). Rather, the formulation of our loss implicitly ensures that the same latent representation, produced by either encoder network, can be understood by either decoder network.

### Extending BABEL to predict protein epitopes

To preprocess single-cell epitope data, we apply a centered log-ratio (CLR) transformation to each cell’s protein counts. This CLR transformation for a count vector *x* measuring *n* proteins in a single cell is given by the following expression, where *g(x)* denotes the geometric mean.

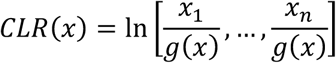

This normalization approach is commonly used for visualizing and modelling epitope data^5,42^.

To extend BABEL to predict these CLR-normalized protein epitopes, we build an auxiliary protein decoder network that takes, as input, BABEL’s sixteen-dimensional latent representation, and predicts CLR-normalized protein counts. This decoder network takes the form of a series of fully connected layers projecting between hidden layers of size 16, 64, 25, and an output layer of 25 dimensions, with PReLU activations applied to each transformation save for the final output layer, which has no activation and directly outputs CLR-normalized values for 25 proteins. Using a similar approach prior works, this protein decoder network is trained using an L1 loss^42^ and paired measurements of protein epitope and RNA expression for single cells. The RNA expression data is first projected into BABEL’s latent representation, which the protein decoder network then uses as input. Note that in this training process, BABEL’s encoder networks and its latent representation are fixed (i.e. their parameters are not updated via backpropagation); the protein network simply uses BABEL’s pre-trained latent representation to make its predictions. Training/validation/test data splits are again defined by scRNA-seq clusters, as was done to train the main BABEL model. To generate protein epitope predictions from scATAC-seq data, we first use BABEL to infer a scRNA-seq profile, and then infer single-cell epitopes from this inferred scRNA-seq profile.

### Evaluation metrics and statistical analysis

We use several metrics to evaluate BABEL’s various outputs. As we consider ATAC chromatin accessibility to be a binary prediction, we use area under the receiver operating characteristic (AUROC) to evaluate the quality of our ATAC predictions. As we consider RNA expression to be a continuous prediction, we use Pearson’s correlation to evaluate the quality of our RNA expression predictions (as well the gene activity score predictions of other tools like ArchR). All such correlations are reported on the log-transformed predicted expression, compared to the log-transformed size-normalized empirical expression. Since scRNA-seq analysis techniques like clustering are predominantly performed on such log-counts, evaluating correlation in log space would be the most relevant metric. Furthermore, all correlations are reported on a random subset of 100,000 entries from the full cell-by-gene matrix. This subsetting is done to reduce the visual clutter of scatterplots, and to ensure the correlations reported are reflective of the scatterplots. AUROC calculations do not use subsetting. For both correlation and AUROC, we effectively consider each gene/peak in each cell a separate observation/prediction.

To generate pseudo-bulk RNA expression signatures that approximate the bulk expression of a tissue, we average the size-normalized, log-scaled expression of every gene across every cell. We then compare these per-gene means across similar samples again using Pearson’s correlation. Pseudo-bulk correlations are not subsetted. However, in cases where we compare bulk expression across two different genome assemblies (i.e. Hg19 and Hg38), the pseudo-bulk is reported on the intersection of genes present in the two assemblies. This is done for the BCC analysis, where we find 17,233 such overlapping genes.

All metrics are calculated using the Python packages Sklearn version 0.21.2^43^, SciPy version 1.2.1^44^, and NumPy. When performing UMAP dimension reduction of single-cell expression data, we apply UMAP projection to the size-normalized, log-scaled expression matrix. UMAP was calculated using functions available through Scanpy^39^ using hyperparameters taken from Seurat’s default settings. We used the package “adjustText” (https://github.com/Phlya/adjustText) to ensure that cluster text labels for these UMAP visualizations did not overlap.

### Analysis on jointly-profiled multi-omic datasets

When evaluating BABEL’s ATAC to RNA predictions on paired ATAC/RNA GM12878 data, or on held-out test clusters, we use the metrics discussed above. We also compare our tool’s predictions to similar outputs generated by ArchR version 0.9.5^21^ and MAESTRO version 1.2.1^22^. We specifically evaluate these methods’ calculations for gene activity scores. Gene activity scores use a weighted sum of each gene’s nearby accessibility to estimate the “activity” or expression of that gene. For ArchR, we use ArchR’s default recommended parameters, as described in the Tutorial (https://www.archrproject.com/articles/Articles/tutorial.html) and online documentation (https://www.archrproject.com/bookdown/index.html), along with the Hg38 assembly. For MAESTRO, we use “enhanced” estimation mode along with default parameters and the Hg38 assembly.

Since we are not aware of any existing tools that can infer ATAC profiles from RNA expression, we are unable to include any reference performance values for this task.

### Unpaired PBMC analysis

Single-cell ATAC-seq healthy PBMC data for analyzing BABEL’s ability to infer scRNA-seq expression profiles from scATAC-seq data is publicly available from 10x Genomics. We used the filtered peak matrix, passed through liftOver for coordinate conversion from Hg19 to Hg38, as input to BABEL. Cell type annotations were derived using scATAC-seq analysis via Signac version 1.0.0, as described here: https://satijalab.org/signac/articles/pbmc_vignette.html. UMAP visualization is generated solely from BABEL’s imputed RNA expression signatures, after size-normalization and log transformation. To contextualize the reconstructed gene expression results, we used scRNA-seq healthy PBMC data publicly available through 10x Genomics, which was then analyzed, clustered, and visualized as described in the corresponding Seurat vignette (https://satijalab.org/seurat/v3.2/pbmc3k_tutorial.html) using Seurat version 3.2.0. All plots showing (predicted or empirical) gene expression are showing size-normalized log expression.

### Basal cell carcinoma analysis

When evaluating BABEL’s ATAC to RNA performance on basal cell carcinoma (BCC) samples, we use scATAC-seq data as published in the original manuscript^10^. As this dataset was originally processed using Hg19, we used the liftOver tool to convert peak coordinates to Hg38 prior to applying BABEL. We use the authors’ publicly available pre-computed Cicero-derived gene activity scores in lieu of calculating them on our own. UMAP plots for this dataset are based on the scATAC-seq measurements; these UMAP coordinates are included in the publicly released dataset as well. Highlighted gene expression is predicted by BABEL based on single-cell ATAC measurements, and is subject to standard size-normalization and log-transformation. Gene activity scores are shown in log scale as well. Pseudo-bulk correlation is reported on the 17,233 genes in the intersection between Hg38 and Hg19 assemblies, as the tissue-matched experiment is analyzed using Hg19^32^.

To find overexpressed marker genes corresponding to a cluster of cells, we perform a Wilcoxon rank-sum test with Benjamini-Hochberg correction, comparing the (predicted) expression of each gene within the cell cluster against the expression of that gene in all other remaining cells. This is done through the functions available in Scanpy. We then take the top 100 most significant results (or fewer if doing so would include non-significant results, i.e. adjusted p > 0.05). The resulting list can then be manually examined for overexpression of key genes.

### Plotting

All plots were generated using Matplotlib^45^, Seaborn (https://seaborn.pydata.org) adjustText (https://github.com/Phlya/adjustText), mpl-scatter-density (https://github.com/astrofrog/mpl-scatter-density), Astropy^46,47^, and Scanpy^39^ libraries under Python 3.7.

## Code availability

All code required to reproduce the BABEL model and our reported results, including data preprocessing and model training, is available at https://github.com/wukevin/babel.

## Data Availability

Human data jointly profiling PBMC cells’ expression and chromatin accessibility is available through 10x Genomics’ <u>data portal</u>. Human data jointly profiling DM and HSR cells’ expression and chromatin accessibility is available through GEO accession GSE160148. SNARE-seq mouse data jointly profiling RNA expression and chromatin accessibility is publicly available at GEO accession <u>GSE126074</u>. PBMC scATAC-seq data is available through 10x Genomics’ <u>data portal</u>. A tissue-matched dataset profiling PBMC scRNA-seq is also available through 10x Genomics’ <u>data portal</u>. Basal cell carcinoma scATAC-seq data is available through GEO accession <u>GSE129785</u>, with gene activity scores available on the original authors’ <u>GitHub page</u>. Tissue-matched basal cell carcinoma scRNA-seq data is available through GEO accession <u>GSE123813</u>.

GM12878 scRNA-seq data is available from GEO accession <u>GSE126321</u>. CITE-seq data on human bone marrow cells is available through GEO accession <u>GSE128639</u>. CITE-seq data on PBMCs is available through the 10x Genomics’ <u>data portal</u>.

## Acknowledgements

We thank the members of the Chang and Zou laboratories for helpful discussions. We thank 10x Genomics for the opportunity to work with data derived from an early, pre-release version of their Chromium Single Cell Multiome ATAC + Gene Expression experimental protocol. K.E.Y. is supported by the National Science Foundation Graduate Research Fellowship Program (NSF DGE-1656518), a Stanford Graduate Fellowship, and a NCI Predoctoral to Postdoctoral Fellow Transition Award (NIH F99CA253729). H.Y.C. is supported by RM1-HG007735 and R35-CA209919. H.Y.C. is an Investigator of the Howard Hughes Medical Institute. J.Z. is supported by NSF CCF 1763191, NIH R21 MD012867-01, NIH P30AG059307, NIH U01MH098953 and grants from the Silicon Valley Foundation and the Chan-Zuckerberg Initiative.

## Author Contributions

H.Y.C. and J.Z. conceived the idea for this project and supervised its execution. K.E.Y. generated data for DM and HSR cell lines. K.E.W. performed data preprocessing, model development, and evaluation of the model, with input from all authors. All authors contributed to analysis and interpretation of results. K.E.W. wrote the manuscript with input from all authors.

## Supplementary Tables and Figures

**Supplementary Table 1:** BABEL performance on test cluster when trained using varying latent dimension sizes. Increasing the size of the latent representation does not consistently improve model performance on the held-out test cluster. Thus, we elect to train BABEL using a 16-dimensional hidden layer to provide the most implicit regularization.

| Test Cluster | Metric | 16-dim | 32-dim | 64-dim |
| --- | --- | --- | --- | --- |
| RNA > RNA | Pearson's $\rho$ | 0.64 | 0.65 | 0.65 |
| ATAC > RNA |  | 0.63 | 0.61 | 0.63 |
| ATAC > ATAC | AUROC | 0.94 | 0.94 | 0.94 |
| RNA > ATAC |  | 0.92 | 0.92 | 0.92 |

**Supplementary Table 2:** Comparison of training independent models (without sharing encoders/decoders) with BABEL on GM12878. The independent models are designed to be architectural copies of the corresponding encoder/decoder combination in BABEL (i.e. the same number of input/output dimensions, number of layers, non-linear activations, etc.), except that the independent models are completely separated and are only trained to perform their singular task. In contrast, BABEL requires that encoders and decoders are interoperable. BABEL outperforms the independent models by a moderate margin for both RNA to RNA and RNA to ATAC translation. The other advantage of BABEL is that due to the interoperability of each subcomponent network, the entire BABEL model contains half of the parameters as the four independent models combined. This makes BABEL faster and easier to train. Overall, this suggests that training interoperable network modules can help BABEL to learn a more efficient, generalizable representation.

| **GM12878** | Metric | BABEL | Independent Models |
| --- | --- | --- | --- |
| RNA > RNA | Pearson's $\rho$ | 0.58 | 0.54 |
| ATAC > RNA |  | 0.58 | 0.59 |
| ATAC > ATAC | AUROC | 0.85 | 0.85 |
| RNA > ATAC |  | 0.66 | 0.63 |

**Supplementary Table 3:** BABEL performance across different held-out test clusters. Each fold represents unique validation and test clusters, cycling through the 5 largest clusters in our data. Performance figures are reported for each fold’s test cluster, which is not used for any training purposes (whereas the validation cluster is used for early stopping and dynamic learning rate adjustments). The intra-domain inference is shown as a sanity check, and is not the primary focus of our work. We observe consistent performance across all folds 2-5; fold 1 exhibits higher ATAC inference performance, along with reduced RNA inference performance. Such variability is expected, as each fold represents evaluation on an entirely different held-out cell type. Notably, this variability does not extend to evaluating BABEL in other contexts, such as GM12878 (Supplementary Table 4) or unpaired PBMC data (Supplementary Table 5). Fold 1 corresponds to the primary model we use throughout the paper, as it is ordinally first and corresponds to using the first and second largest clusters as validation/test.

| Test Cluster | Metric | Fold 1 | Fold 2 | Fold 3 | Fold 4 | Fold 5 |
| --- | --- | --- | --- | --- | --- | --- |
| RNA > RNA | Pearson's $\rho$ | 0.64 | 0.82 | 0.81 | 0.84 | 0.84 |
| ATAC > RNA |  | 0.63 | 0.79 | 0.78 | 0.81 | 0.82 |
| ATAC > ATAC | AUROC | 0.94 | 0.90 | 0.90 | 0.91 | 0.91 |
| RNA > ATAC |  | 0.92 | 0.86 | 0.86 | 0.88 | 0.87 |

**Supplementary Table 4:** BABEL performance on the GM12878 paired ATAC/RNA data. These folds correspond to the same data splits as used in Supplementary Table 3. Performance is always shown on the same set of GM12878 cells, but the model itself changes as its training and validation sets vary. We observe very little variation in performance, which indicates that BABEL is robust to variation in its training set and exhibits similarly strong generalizability regardless.

| GM12878 | Metric | Fold 1 | Fold 2 | Fold 3 | Fold 4 | Fold 5 |
| --- | --- | --- | --- | --- | --- | --- |
| RNA > RNA | Pearson's $\rho$ | 0.58 | 0.60 | 0.58 | 0.56 | 0.58 |
| ATAC > RNA |  | 0.58 | 0.56 | 0.53 | 0.55 | 0.58 |
| ATAC > ATAC | AUROC | 0.85 | 0.85 | 0.84 | 0.84 | 0.85 |
| RNA > ATAC |  | 0.66 | 0.67 | 0.69 | 0.65 | 0.56 |

**Supplementary Table 5:** BABEL’s pseudo-bulk concordance on unpaired PBMC cells across cross-validation folds. As in Supplementary Table 4, each fold here represents a different model that was trained using a different training and validation set spanning DM, HSR, and PBMC cells. We use pseudo-bulk correlation here, which measures the concordance in the average expression per gene across cells, as this dataset is not paired. Despite variations in the training set, BABEL exhibits uniformly high performance across all folds. This further indicates that BABEL is a robust model that consistently learns generalizable patterns.

| Unpaired PBMC performance | Fold 1 | Fold 2 | Fold 3 | Fold 4 | Fold 5 |
| --- | --- | --- | --- | --- | --- |
| Pseudo-bulk Pearson's $\rho$ | 0.89 | 0.89 | 0.91 | 0.91 | 0.89 |

**Supplementary Figure 1:**
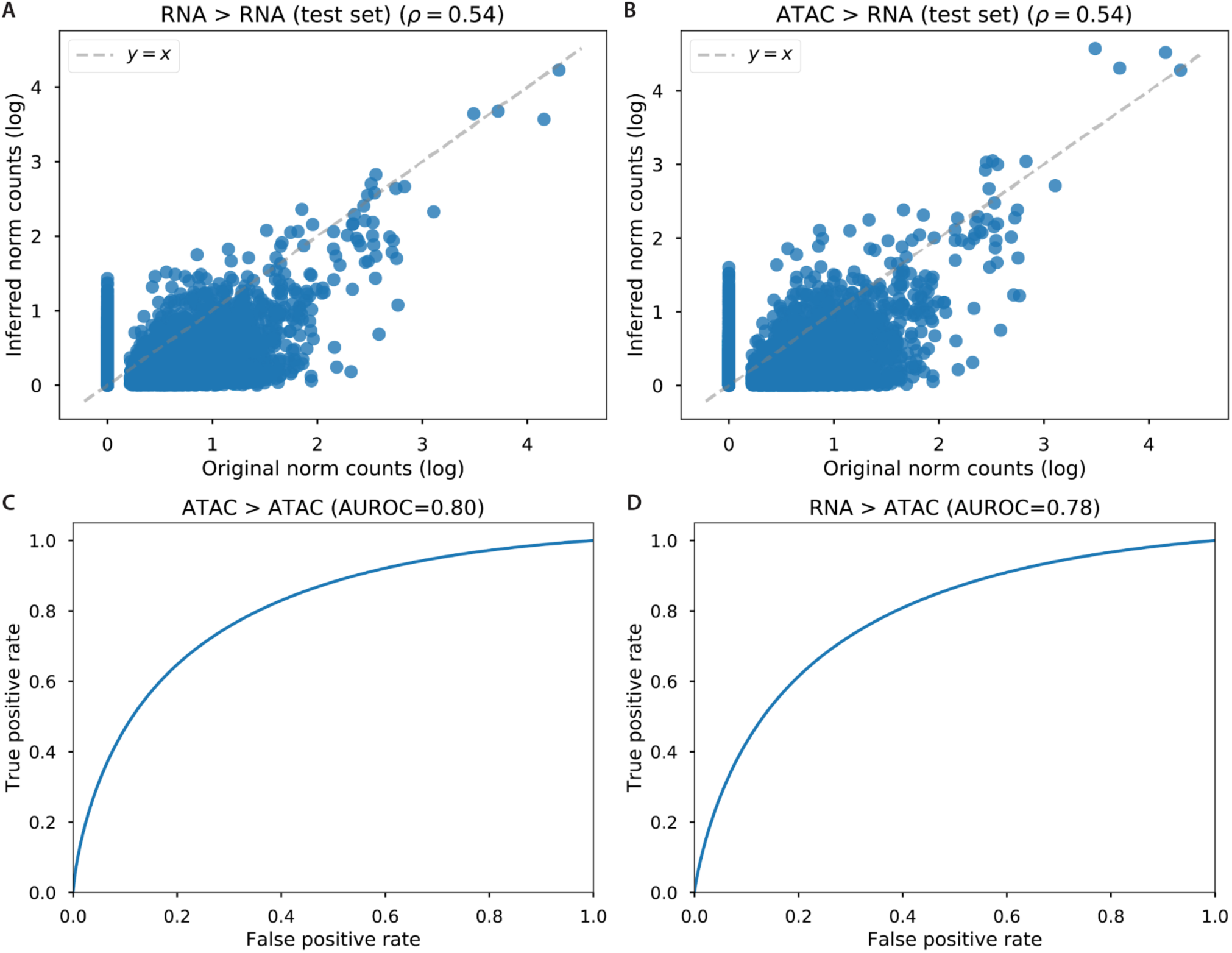
BABEL’s performance when trained and tested on the SNARE-seq mouse data. Test cluster consists of 1308 cells. (A) Intra-domain RNA to RNA inference performance. Each point represents the expression of one gene in one cell (100,000 such entries are randomly chosen for visual clarity) within the held-out test cluster; the x-axis represents the empirically measured expression of that gene in that cell, while the y-axis represents the inferred expression. (B) Cross-domain ATAC to RNA inference performance, formatted as in panel (A). (C) ATAC to ATAC intra-domain inference. (D) RNA to ATAC inference performance. We observe that all four possible encoder/decoder combinations perform well, indicating that BABEL can be successfully trained on non-human data generated using different experimental protocols.

**Supplementary Figure 2:**
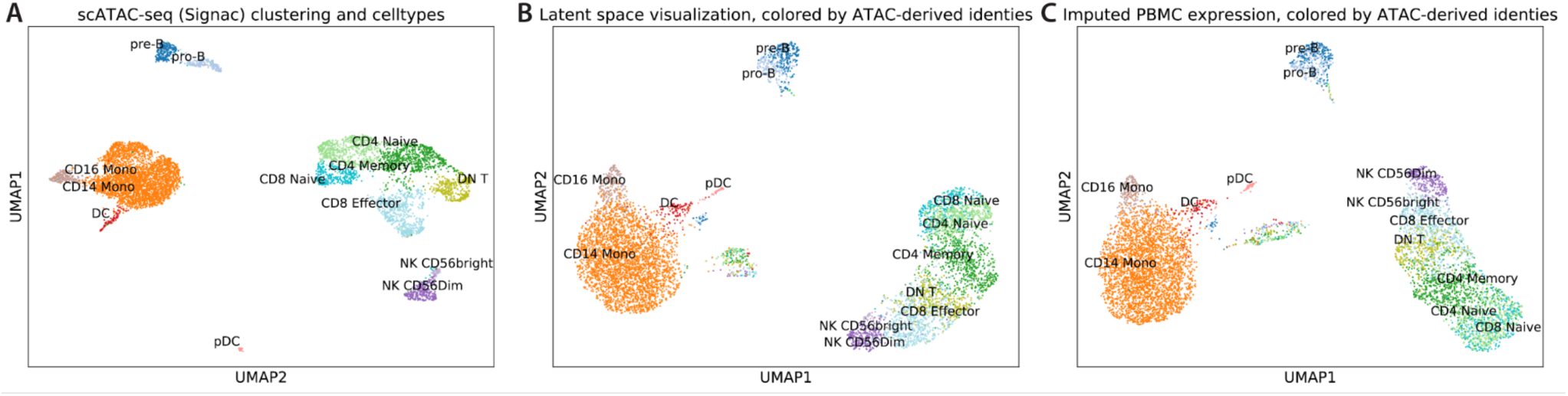
Cell type continuity between the three main steps within BABEL: input (A), latent representation (B), and output (B). All panels show cells colored by their ATAC-based identities to clearly demonstrate how these are tracked through BABEL. (A) UMAP visualization of the input scATAC-seq human PBMC data, generated via the Signac tool. Axes are swapped for easier visualization with other panels. We see separation between a primarily monocyte cluster (left), a B cell cluster (top), and a CD4/CD8/NK cluster (right). These three main clusters are retained after BABEL projects this scATAC-seq data into its latent representation, which is visualized in panel (B). This is strong evidence that BABEL’s latent space has learned the relationships between cell types, despite having no prior information about what these cell types are. This latent representation is then used to infer the scRNA-seq expression profiles using the RNA decoder, the output of which is visualized in (C). This panel is a reproduction of Figure 3A and is very similar to an empirical scRNA-seq experiment done on the same tissue (Figure 3B), and is included here for ease of comparison. We see that throughout all stages in BABEL’s ATAC to RNA translation, BABEL preserves the overall biological relationships between these cells.

**Supplementary Figure 3:**
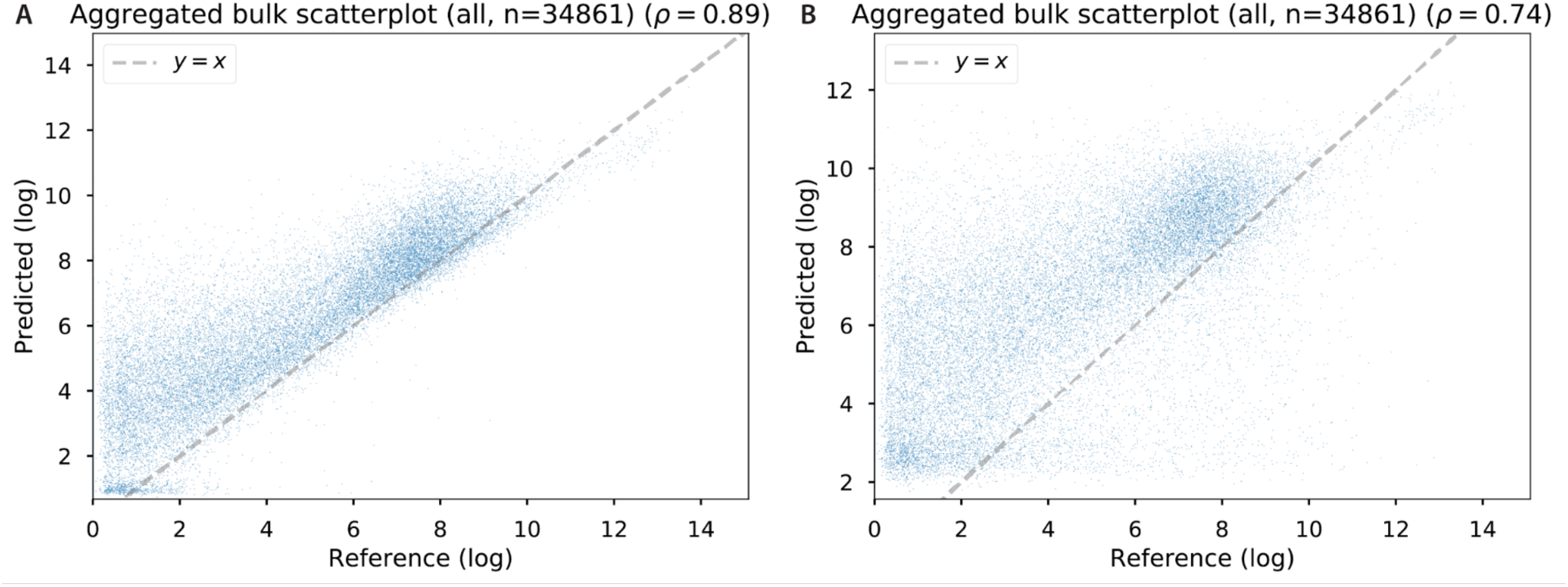
Pseudo-bulk correlation of different versions of BABEL on PBMC samples. In both plots, the x-axis represents measured scRNA-seq expression on PBMC data, and the y-axis represents inferred single-cell expression from scATAC-seq on PBMC data. Since the two modalities are not paired, but are from the same tissue, we compute the average expression per gene across all cells from each – the pseudo-bulk expression. Due to the large number of genes (n=34,861), we plot these as density heatmaps showing log-scaled counts. (A) Shows pseudo-bulk expression concordance when BABEL is trained on DM, HSR and PBMC data, and (B) shows pseudo-bulk expression concordance when BABEL is trained on just DM and HSR cells, excluding PBMCs. Including PBMCs in the training data improves pseudo-bulk expression correlation with the measured scRNA-seq. This demonstrates the utility of pseudo-bulk correlation in allowing us to objectively quantify the trustworthiness of BABEL’s single-cell predictions even without paired ground truth measurements.

**Supplementary Figure 4:**
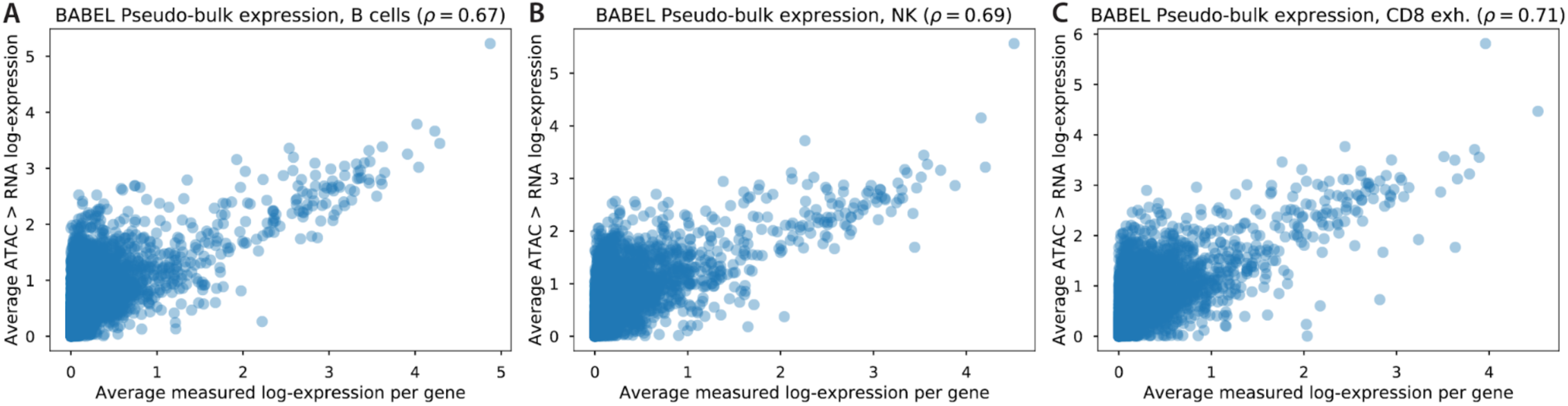
Pseudo-bulk correlation of BABEL on BCC samples, broken down by various cell types. X-axis represents expression within a tissue-matched scRNA-seq study averaged across cells; y-axis represents BABEL’s single-cell ATAC to RNA predictions, also averaged across cells. Each panel shows pseudo-bulk expression averaged across a different cell type within the global set of cells: B cells (A), natural killer cells (B), and CD8 exhausted T cells (C). We find that the pseudo-bulk correlation is consistent for each cell type, not just for the global population of cells (FIgure 4A). This provides further evidence that BABEL produces high-quality predictions that are not simply general bulk estimates.

**Supplementary Figure 5:**
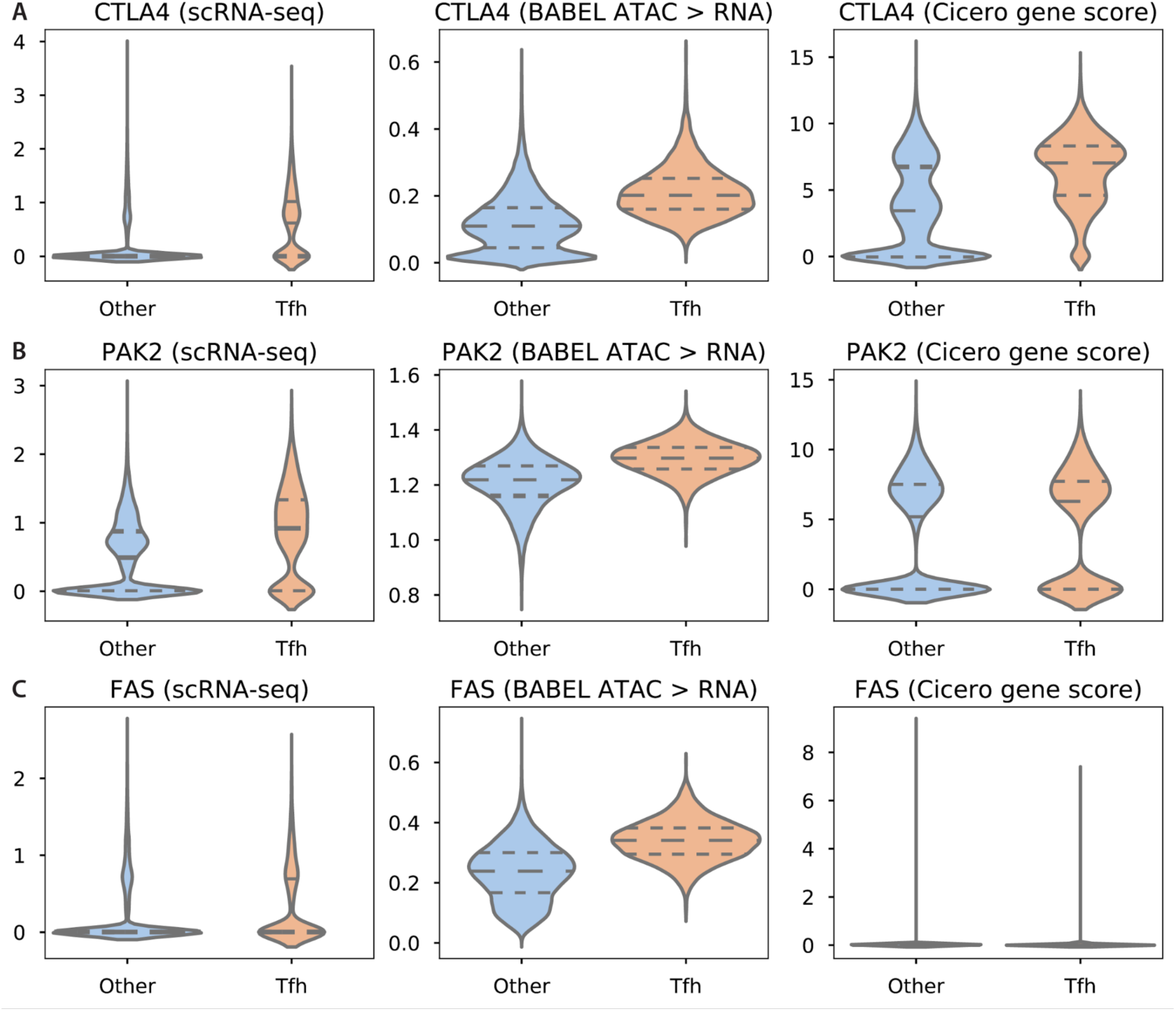
Predicted versus empirical overexpression of immunosuppressive genes in Tfh cells. Each gene shown in each row of plots was predicted to be significantly overexpressed by BABEL. All violin plots show expression in log scale, with dashed horizontal lines indicating upper/lower quartiles and median. Since scRNA-seq, BABEL, and Cicero are all normalized differently, absolute numeric values are not comparable. (A) Shows the expression of *CTLA4* within the Tfh cell cluster, compared with remaining cells, as measured by a tissue-matched scRNA-seq experiment (left), as predicted from scATAC-seq data by BABEL (center), and as inferred by Cicero’s gene activity scores (right). In this case, both BABEL and Cicero predict overexpression, which is validated by the experimental data (p < 0.05, Mann-Whitney test). (B) Similar comparison for *PAK2*, where we see very strong overexpression in Tfh cells in empirical scRNA-seq (left panel). BABEL (center) more strongly predicts overexpression here compared to Cicero (right), though both are significant (p < 0.05, Mann-Whitney test). (C) Comparison for *FAS*, which exhibits subtle overexpression in scRNA-seq (p < 0.05, Mann-Whitney test). Here, BABEL is able to correctly predict a slight overexpression in the Tfh cluster (p < 0.05, Mann-Whitney test), whereas gene activity scores predict no statistically significant overexpression (p > 0.05, Mann-Whitney test).

**Supplementary Figure 6:**
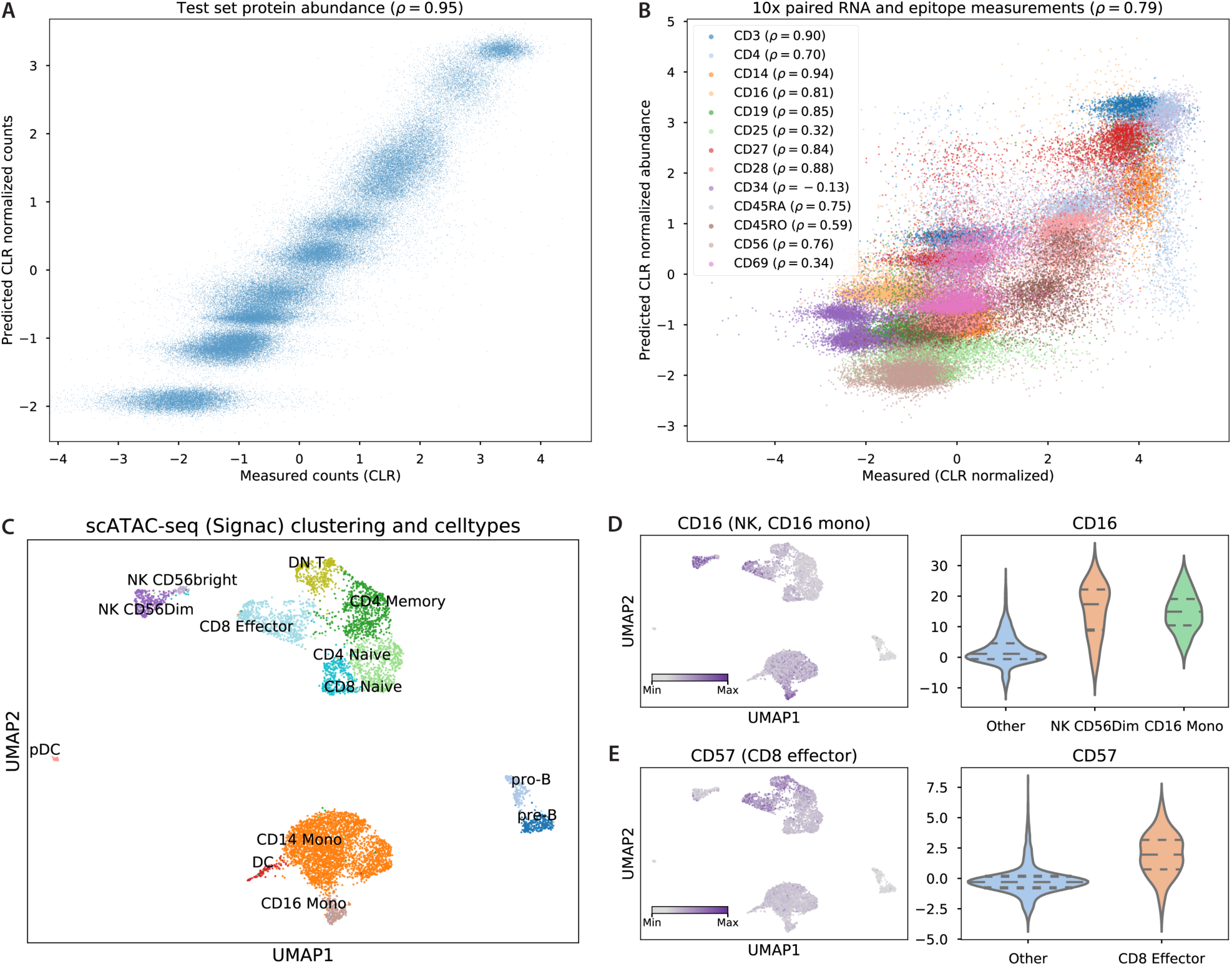
Extending BABEL to infer single-cell protein epitopes. Using a set of bone marrow CITE-seq measurements, we train a protein decoder network that takes BABEL’s pre-trained latent representation and predicts CLR-normalized protein counts. (A) Shows performance of this approach on a held-out test set of 4,032 bone marrow cells, and (B) shows predicted protein expression for 5,221 PBMC cells (also profiled via CITE-seq), colored by protein. Only proteins present in both the bone marrow and PBMC CITE-seq measurements are shown. In both panels, the x-axis denotes the CLR-normalized measured protein abundance, the y-axis denotes the protein decoder’s predictions given BABEL’s pre-trained latent representation based on these cells’ RNA signatures, and each point represents a single cell’s abundance of a given protein. In both cases, we see that BABEL can be successfully extended to predict epitope measurements from scRNA-seq measurements (Pearson’s correlations of 0.95 and 0.79, respectively). (C-E) show that we can use this protein decoder to predict epitopes from scATAC-seq, even without training this specific modality pair. (C) Visualizes a PBMC scATAC-seq dataset (same dataset as Supplementary Figure 2A). We use these chromatin accessibility profiles to impute epitopes, shown in (D, E). (D) highlights predicted abundance of CD16 for each cell. CD16 is a marker associated with CD16 monocytes and CD56Dim natural killer cells^20^ – a pattern that is recapitulated by BABEL’s ATAC to protein predictions (violin plot, right panel). (E) shows predictions for CD57, which BABEL correctly predicts to be most abundant in CD8 effector cells.

## Supplementary Note

### Chromosome-aware ATAC encoder/decoder architecture

BABEL’s component networks that encode and decode ATAC chromatin accessibility data are designed to leverage the intuition that most chromatin interactions occur on an intra-chromosomal level^28^. Accordingly, we prune many of the inter-chromosomal parameters in BABEL’s ATAC encoder and decoder networks. In the following, we provide an estimate for how many parameters this approach saves. We describe the ATAC decoder, but the analysis is the same for the encoder.

For simplicity, we only consider parameters directly involved in linear matrix transformations, disregarding parameters needed for activation functions, batch normalization, etc. When trained on human data, BABEL learns to predict 223,897 peaks across 22 autosomes; we simplify this as 10,000 peaks per chromosome. Recall that the shared latent space is 16-dimensional, and maps to a concatenated layer of 16 dimensions per chromosome. Thus, the latent to “concatenated” layer contains 16 x (22 x 16) = 5,632 parameters; this is constant whether we are using chromosome-specific architectures or not, and allows for limited representation of inter-chromosomal interactions in the former case. Under the chromosome-aware design, this then leads to 22 chromosome sub-units, mapping from 16 to 32 to 10,000 output dimensions: 22 x (16 x 32 + 32 x 10000) = 7,051,264 weights. Under the chromosome-agnostic design, we would instead have (22 x 16) x (22 x 32) + (22 x 32) x 220000 = 155,127,808 weights, or 22 times the number of weights for the final two layers.

### BABEL’s composition of encoder and decoder networks and their shared latent representation

BABEL takes the approach of composing two encoder and two decoder neural networks, such that the encoder and decoder networks are interoperable. We benchmarked this approach against a naive approach that learns four completely separate neural networks with no interoperability constraints. These four neural networks are architecturally identical to each of BABEL’s four encoder/decoder combinations, but have no interoperability requirements. Compared to BABEL, this naive approach uses twice the number of overall parameters, as it requires 4 separate pairs of decoders and encoders, compared to BABEL’s combinatorial use of 2 encoders and 2 decoders.

We performed our evaluation using the GM12878 paired data, as this dataset is fully external to model training and is thus the most stringent measure of model generalizability and performance. We found that for three of the four tasks, BABEL either matches or outperforms these independent models (Supplementary Table 2). This suggests that our interoperability constraint has no overall negative impact on performance, and seems to actually be helping BABEL learn more general representations and functions for translating cellular modalities. Furthermore, these results confirm the efficacy of our enforcement of a shared latent representation via loss-induced encoder-decoder interoperability. In contrast, prior works translating between multi-omic profiles did not leverage paired measurements, and consequently required complex adversarial networks or expensive manifold alignment strategies to align points in a similar shared latent representation ^15–17^. Since BABEL’s latent representation is already aligned (otherwise BABEL’s performance wouldn’t be able to match/exceed that of single-purpose dedicated models), such manifold alignments approaches would only add unnecessary complexity.

Beyond being a well aligned shared representation, BABEL’s latent representation serves as a critical information bottleneck as well. Namely, by restricting the size and thus possible information content of the latent representation that all encoder/decoder combinations must use, we restrict our model to focusing on the most important factors driving cell-to-cell variation. This provides implicit regularization and is similar to how tools like principal component analysis can denoise data by compressing data into the top few principal components. When designing BABEL, we considered several sizes for this latent representation. We found that increasing the dimensionality of the latent space to 32 or even 64 (compared to 16) provided no consistent benefit in BABEL’s ability to perform well on the test set (Supplementary Table 1). In the absence of any meaningful performance impact, we elected to use the most restrictive latent representation of 16 dimensions to minimize potential model overfit.

Furthermore, we see evidence that BABEL’s 16-dimensional latent space encodes biologically meaningful relationships. If we apply BABEL’s ATAC encoder network to scATAC-seq PBMC data and visualize the resultant latent representation using UMAP, coloring each point by its cell type, the resulting visualization (Supplementary Figure 2B) is highly similar to reference plots generated by applying UMAP to empirical scATAC-seq (Supplementary Figure 2A) and scRNA-seq (Figure 3B). Remarkably, not only are cells of the same cell type grouped together in the latent representation, cells of similar cell types appear to be closer in this latent space as well. For example, CD14 and CD16 monocytes occupy similar (but distinct) regions in the latent space, which is reflective of their biological similarity compared to other PBMC cell types; the same can be said of pro-B/pre-B cells, natural killer subtypes, etc. This suggests that BABEL’s latent space effectively captures important biological variation, even without being given prior information regarding cell type identities or relationships. In fact, we suspect that this “biologically continuous” representation improves generalization, as it allows for efficient interpolated representation of a spectrum of intermediate cell types within the latent space. This is especially evident compared to a hypothetical alternative where the latent space naively represents each cell type independently of other cell types; this would necessitate a great amount of training data to learn every individual cell type via rote memorization and would likely struggle to generalize to even slight variations on previous data.

### Understanding BABEL’s performance on PBMCs through data ablation

We hypothesize that one of the key factors driving BABEL accuracy in predicting PBMC cells’ single-cell gene expression is the inclusion of PBMCs in its training data. To quantitatively measure this in the absence of paired measurements, we compare pseudo-bulk expression signatures (i.e. the mean across cells of every gene’s expression). We observe that BABEL’s imputed pseudo-bulk PBMC expression exhibits a Pearson’s correlation of 0.89 when compared to a pseudo-bulk derived from the aforementioned PBMC scRNA-seq experiment (Supplementary Figure 3A). This correlation is highly consistent across different versions of BABEL trained using different cross-validation folds (Supplementary Table 5). However, if we remove PBMCs from BABEL’s training set and re-train BABEL from scratch, this pseudo-bulk correlation drops to 0.74 (Supplementary Figure 3B). This analysis concretely demonstrates the benefits of training BABEL on a similar group of cells, and also highlights how aggregate metrics may be used to quantify BABEL’s performance in a general setting without paired measurements.

